# Robust Deep Learning-based 3D Segmentation and Morphological Analysis of Mitochondria using Soft X-ray Tomography

**DOI:** 10.1101/2025.10.16.682919

**Authors:** Arun Yadav, Anshu Singh, Aneesh Deshmukh, Pushkar Bharadwaj, Anuj Baliyan, Kate White, Jitin Singla

## Abstract

Mitochondrial morphology is crucial for cellular function, but large-scale analysis is limited by challenges in high-resolution imaging and segmentation. MitoXRNet, a compact 3D deep-learning model, efficiently segments mitochondria and nuclei from Soft X-ray Tomography data using multi-axis slicing, Sobel-based boundary enhancement, and combined BCE–Robust Dice loss. With 1.4M parameters, it achieves a Dice score of 73.8% on INS-1E cells, outperforming existing models. Automated analysis indicated that glucose induced larger mitochondria and higher matrix density, and that GIP and GKA induced smaller and denser mitochondria, highlighting previously unreported β-cell mitochondrial remodeling. MitoXRNet allows for scalable profiling of organelle-level morpho-biophysical data.

**Highlights:**

- A data-efficient method for 3D segmentation of mitochondria and nucleus from Soft X-ray tomograms.
- Incorporates domain-specific Sobel filter-based preprocessing to improve segmentation accuracy and quality under imperfect or noisy labels.
- Enables rapid and automated analysis of mitochondrial morphology, facilitating quantitative assessment of pharmacological effects on cellular ultrastructure.

**Graphical Abstract:** 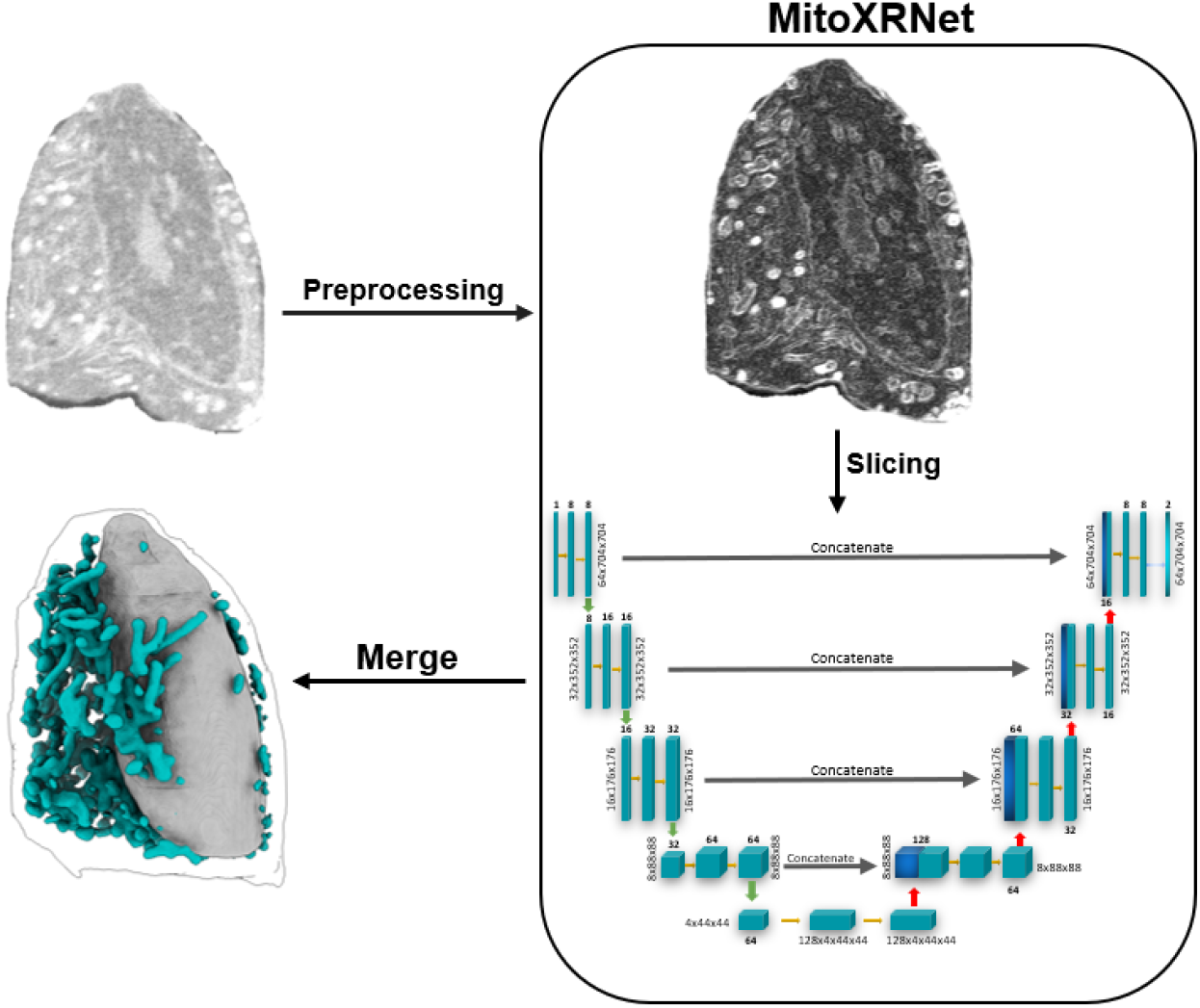

## 1. Introduction

Inter-organelle interactions, organelle distribution, shape and size play a vital role in normal cellular function. Mitochondria among all organelles is an important marker to study the cellular health and stress state. There are detailed studies showing close structure-function relationship between cellular and mitochondrial health. Mitochondria exhibit distinct morphology under stressed and healthy state, including change in spatial distribution, fission-fusion dynamics and metabolic state. The detailed quantification of these morphological features can help understand the structural state of whole mitochondrial network inside cells, which can emerge as an important structural biomarker to measure the stress. Rapid and automated quantification techniques can facilitate in evaluating the effect of pharmacological interventions in reducing the cell stress and study the shift of cellular state from diseased to more healthy state.

Quantifying the structural organization of organelles, especially mitochondria, is challenging due to their complex architecture, the wide range of spatial scales in cell biology, and the limitations of traditional imaging methods such as fluorescence microscopy, which requires probe tagging and disturbs the native state, or electron microscopy, which restricts the imaging window or requires sectioning that limits data collection. Among available imaging techniques, Soft X-ray Tomography (SXT) can capture the high-resolution images of the whole 3D cells in their native state (without staining) and within a span of 5 min ^1–3^. SXT provides an opportunity to study cells at ultrastructural (mesoscale) scale. It allows us to study cellular organelles, their morphology and spatial alignment within the cell (Elgass et al., 2015; Loconte et al., 2022; White et al., 2020). The systematic study of mitochondrial morphology, temporal dynamics and the effect of drug treatments require binary masks of mitochondria segmented out from noisy grayscale 3D SXT tomograms. Most of the analysis currently available in the literature has been conducted on manually segmented masks. Manual segmentation is a time consuming and error prone process. It is also prone to human variability. An expert can take up to 2 days to manually segment complete mitochondrial network per tomogram. This drastically limits the potential of SXT for quantifying morphological changes which require hundreds of segmented cells.

Deep Learning (DL) based segmentation can play an important role in providing efficient solutions for segmenting cellular organelles from tomograms. Although a substantial amount of literature has been published around cell and mitochondria segmentation using various imaging modalities, such as fluorescent microscopy (Fischer et al., 2020; Hultgren et al., 2024; Ounkomol et al., 2018; Padovani et al., 2022; Stringer et al., 2021; Zargari et al., 2024) and electron microscopy (Conrad & Narayan, 2022; Heinrich et al., 2021; Jang et al., 2025; Shi et al., 2024; Taşel & Çi□ftci□, 2025), etc., limited studies are available for Soft X-ray Tomography focusing on either cytoplasm, nucleus, vacuoles, with poor performance on mitochondrial segmentation (Chen et al., 2024; Chueh et al., 2025; Erozan et al., 2024, 2025; Francis et al., 2020; Li et al., 2022). Electron microscopy generates high resolution images and has sharp organelle boundaries, whereas SXT tomograms have low contrast, making it challenging to segment cellular structures. On other hand, fluorescence images do not require manual segmentation, as the ground truth can be generated by thresholding the fluorescent labels, with little to no human error. Also, collecting large amounts of training data is relatively more feasible.

DL-models around soft X-ray tomography are comparatively less explored. Firstly, because the contrast overlap between organelles makes it challenging, secondly, limited manually labelled data is available. (Francis et al., 2020) proposed a CNN-based method to apply partially labeled data for full label segmentation and showed that mixed labels (partially and fully annotated) perform better than standard fully annotated data-based training (Francis et al., 2020). Authors achieved a 57.5% dice score on the mitochondrial network. Further, (Li et al., 2022), published a method wherein they trained 2D U-Net architecture on slices of the tomograms for segmenting cytoplasm, nucleus, mitochondria and vesicles. Using 2D slices from all three-axis generated ample amount of training data from the limited 3D annotated data. For mitochondrial segmentation, they achieved a dice score of 68.34% on three in-domain test cells. Two more studies focusing solely on cytoplasm (Erozan et al., 2024) (ACSeg) and nucleus (Erozan et al., 2025) segmentation have also been recently published. Additionally, models like nnU-Net (Isensee et al., 2021) self-configures U-Net architecture, loss functions and preprocessing techniques based on input data, making them invariant to imaging techniques. It has been shown to work well on large numbers of biomedical imaging data for segmentation. We hypothesize that organelle segmentation requires 3D context and specialized preprocessing techniques to address the low contrast between organelles and that previously used 2D slicing may not preserve the full context of organelles and subcellar structures. A model can easily confuse the circular cross-section of mitochondria and vesicles without any volumetric context.

In this paper we propose MitoXRNet: a data efficient 3D U-Net based auto-segmentation technique to segment mitochondria from whole-cell SXT tomograms. The proposed method uses overlapping 3D slices of the whole tomogram along the three axes for generating large training data from limited annotated dataset. We applied a Sobel filter–based custom preprocessing pipeline to highlight the mitochondria boundary. We trained and tested the model with two loss functions: the conventional Binary Cross-Entropy (BCE) loss and a combined BCE with Robust Dice loss. BCE focuses on voxel-level accuracy, while the Robust Dice component improves the overall overlap between predictions and ground truth. In addition, Robust Dice effectively handles class imbalance, which is critical in our case since the mitochondrial volume is substantially smaller than the cell volume.

We trained and tested our method on the same INS-1E (pancreatic β-cell line) SXT dataset as done by (Li et al., 2022), along with additional cells in the training data set. We also used another independent dataset treated with various drugs, collected from the same source but with different set of microscope calibration and settings (better zone plate resolution). This separate dataset was used to measure out-of-domain performance, evaluating the generalizability and robustness of the models.

We trained models with two different depths and varying kernel filters:

i. A smaller model with only 1.4 million (M) parameters outperformed (Li et al., 2022) results and state-of-the-art models like nnU-Net (Isensee et al., 2021) (auto configured with 31.2M parameters) on the in-domain test dataset. The smaller model could not generalize well compared to nnU-Net on out-of-domain dataset.
ii. A larger model with 22.6M parameters that showed similar performance on in-domain dataset and outperformed nnU-Net on out-domain test dataset.

MitoXRNet results showed improved mitochondrial segmentation with adequate preprocessing and proposed 3D slicing. Nucleus was also segmented along with mitochondria to provide an anchor for any downstream analysis of spatial distribution in the cytosolic region of the cell. We also showed the case study of automated segmentation in quick quantification of mitochondrial morphology revealing increased level of fragmentation and increased metabolic activity in cells treated with glucose + glucokinase activator (GKA-50) and glucose + gastric inhibitory polypeptide (GIP). Glucose increased mitochondrial volume and matrix density (voxel LAC), while GIP and GKA increased mitochondria number, reduced volume, and elevated density, indicating smaller, denser, and more dynamic mitochondrial populations. These results reveal unreported effects of GIP and GKA on β-cell mitochondrial architecture.

With the increasing availability of SXT (Harkiolaki et al., 2018) and its use in studying cellular architecture and morphological changes under various stress and disease conditions, the requirement for auto-segmentation tools will only increase. SXT is also a growing imaging modality with potential to contribute to whole-cell modeling efforts (Loconte et al., 2023; Singla et al., 2018; Singla & White, 2021). Our method can help researchers segment hundreds of tomograms with much higher efficiency and study architectural changes to mitochondria using soft X-Ray tomography.

## 2. Materials and Methods

### 2.1 Soft-X-ray tomography

Image projections were collected at 517 eV using XM-2 (Le Gros et al., 2014) soft X-ray microscope at the National Center for X-Ray Tomography at the Advanced Light Source (Lawrence Berkeley National Laboratory); the microscope was equipped with a 50-nm zone plate. During data collection, cells were maintained in a stream of helium gas cooled to liquid nitrogen temperature. Projection images were collected sequentially around a rotation axis of 180^*o*^(capillary axis), with 2^*o*^ increments and an exposure time of 350ms. Tomograms were reconstructed from the acquired projections using AREC3D, with voxel intensity normalized to attribute accurate linear absorption coefficient (LAC) values across all samples.

### 2.2 Dataset

The first dataset was obtained from previous publications by (White et al., 2020) and (Loconte et al., 2022), containing a total of 55 tomograms of Rat insulinoma INS-1E cell line from Addex Bio, collected with 60-nm zone plate. The second dataset used for out-of-domain testing was obtained from Deshmukh et al.(Deshmukh et al., 2025), containing 44 tomograms of Rat insulinoma INS-1E cells from Pierre Maechler’s lab (University of Geneva) (Merglen et al. 2004), collected with 50-nm zone plate.

### 2.3 Cell, Nucleus and Mitochondria masks labels

The ground truth cell masks were generated using ACSeg Click or tap here to enter text.(Erozan et al., 2024). The cell masks from ACSeg were refined manually by the experimentalist to prevent any error accumulation in the training of mitochondria and nucleus segmentation. The ground truth masks for mitochondria and nucleus were manually segmented using Amira (FEI) software.

### 2.4 Preprocessing of 3D Soft X-Ray tomograms

To enhance mitochondrial structures in the 3D soft X-ray tomograms, we applied a combination of Gaussian smoothing and Sobel-based high-pass filtering. First, the raw image volume *I* was smoothed with a Gaussian filter *G*_*σ*_ of standard deviation *σ*=2, generating

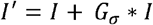

where * denotes convolution. Edge information was then extracted using a 3D Sobel operator along the *x, y* and *z* directions:

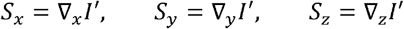

and the Sobel magnitude was computed as

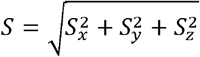

This high-pass response was combined with the original image in a nonlinear form:

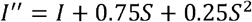

Where the quadratic term enhances strong edges while suppressing noise. Finally, both *I*″ and *I* were normalized to the range [0,1] and multiplicatively combined to yield the preprocessed tomogram:

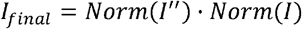

This procedure emphasizes fine mitochondrial boundaries while retaining overall structural contrast, thereby facilitating more accurate segmentation.

### 2.5 Resizing and 3D Slicing

Soft X-ray tomograms have different dimensions along each axis and between different datasets due to different resolutions and variable field of view. The tomograms after preprocessing and cytoplasm segmentation using ACseg (Erozan et al., 2024), were resized to 704 × 704 × 704 voxels using zero center padding. Further, the tomograms were sliced along the three axes independently, generating 3D slices of size 64 × 704 × 704, with a sliding window of 32 voxels. This creates an overlap of 32 voxel depth between two 3D slices. Single tomogram generates 21 3D slices in each direction with a total of 63 3D slices.

### 2.6 Model training

For training the model out of 55 tomograms in first dataset, 49 tomograms were used for training, 3 for validation and 3 for testing. Another dataset with 44 tomograms was used for out-of-domain testing. In total we used 3087 training, 189 validation and 189 test slices from the 55 padded tomograms.

In this work, we implemented two configurations of 3D U-Net based on number of layers and filters. One model with 1.4M parameters using 4 encoder-decoder blocks along with Batch Normalization(Ioffe & Szegedy, 2015), with 1, 8, 16, 32 and 64 (bottleneck Conv layer) kernels in each block respectively. Another model with 22.6M parameters using 5 encoder-decoder blocks and 1, 16, 32, 64, 128 and 256 (bottleneck Conv layer) kernels. Both the models were trained with a batch size of 4 slices using Adam optimizer(Kingma & Ba, 2014) with a fixed learning rate of 1e^-4^. Both the models were trained under two loss functions independently, corresponding to two separate training regimes.

### 2.7 Loss functions

This study uses two loss functions. The formulation of both loss function over a single 3D-slice is defined below.

1. Binary Cross-Entropy (BCE) Loss

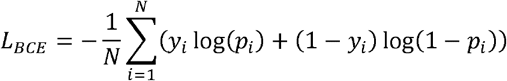

where *N* is the total number of voxels in the slice, *y*_*i*_ is the true binary label and *p*_*i*_ = σ(*z*_*i*_) is the predicted probability at voxel *i* computed using sigmoid function.
2. Combined Loss of Robust Dice and BCE Loss

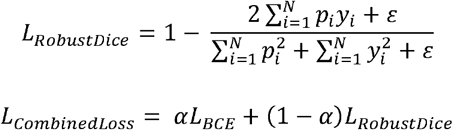

where, *N, y*_*i*_ and *p*_*i*_ are as defined above, *ε* is a small value for numerical stability and α is the hyperparameter for linear combination of BCE and Robust Dice loss. In this study, we used *α* = 0.5.

### 2.8 Merge 3D slice predictions

The training of the model occurs at a single 3D slice level, however, to obtain the segmented mask for complete mitochondria and nucleus. Predictions from all overlapping 3D slices along the three axes are combined into a single mask.

For each slice *s*, the model outputs logits for two classes (i.e. nucleus and mitochondria). Let 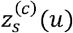 be the logit predicted for class *c* ∈ {*nucleus, mitochondria*} at local coordinate *u* in slice *s*. These logits are converted to probabilities independently for each class using the sigmoid functions:

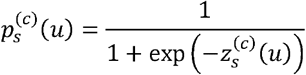

Once probabilities are computed from all 63 3D-slices (21 along each axis), they are mapped back to the global 3D volume of the cell. Because the same voxel is predicted multiple times (due to slice overlaps and predictions from all three axes), voxel-wise probabilities are averaged across all occurrences for each class independently. This generates two probabilistic masks, one for each class. Next, class-wise probabilistic masks are merged into a single two-class probabilistic mask by assigning each voxel to the class with higher probability. Finally, to obtain binary segmentation masks, a threshold is applied. Voxels with probability below the threshold are classified as background, while remaining voxels are assigned to either nucleus or mitochondria based on higher probability.

### 2.9 Computing Dice Score

The dice score (Dice similarity Coefficient) is used to quantify the overlap between predicted and ground truth segmentation masks for each class (organelle). The Dice score ranges from 0 to 1, where the higher value indicates the higher agreement between predicted and ground truth masks. Dice score is computed for a whole 3D organelle mask after merging the probabilistic predictions of each slice and thresholding the probability into binary prediction. It is defined as:

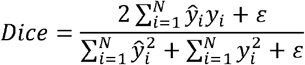

where *N* is the total number of voxels in tomogram, *y*_*i*_ is the true binary label and *ŷ*_*i*_ = (*p*_*i*_ ≥ *threshold*) is the predicted binary mask at voxel *i* computed by thresholding the probability mask.

### 2.10 Computing Individual mitochondria volume and count

The post-segmentation mitochondrial mask is a single binary mask in which all mitochondrial voxels were assigned to the same label. To quantify individual mitochondria, this binary mask was processed using the *label* function from the *scipy*.*ndimage* module (version 1.10.1) (Virtanen et al., 2020). A 3D connectivity structure (26-connected neighborhood) was used to ensure that all spatially contiguous mitochondrial voxels were grouped as a single object. The labeling step produced a unique integer index for each disconnected mitochondrial component, allowing us to compute the total number of mitochondria. The volume of individual mitochondrion was obtained by voxel count associated with its corresponding label and multiplying by the physical voxel volume.

### 2.11 Statistical analysis and data representation

Statistical analysis was performed to evaluate the significance of differences in mitochondrial morphology and molecular density across experimental conditions. All analyses were carried out using the Kruskal–Wallis test, followed by Dunn’s post hoc multiple comparison test for intergroup analysis using python package (*statsmodels* (version 0.13.5) and *scikit_posthoc* (version 0.9.0)). Since each condition contained many mitochondria, results are summarized using boxplots that depict the overall distribution, mean, median, and quartile range for each group. Red asterisks without connecting brackets indicate statistical significance when compared with the control condition (No Stimulation), whereas red asterisks accompanied by brackets denote significant differences between the corresponding treatment groups. Levels of significance are represented *p<0.05; **p<0.01; ***p<0.001, ****p<0.0001.

## 3. Results

### 3.1 Auto-segmentation workflow

A single SXT tomogram contains whole 3D cell(s) inside a glass capillary, typically oriented vertically. To develop an accurate segmentation of mitochondria and nucleus, first single cells were segmented out using ACSeg (Erozan et al., 2024), available on Biomedisa. The cell masks obtained from ACSeg were manually corrected to limit the error propagation in the downstream training process. The rest of the workflow uses tomograms masked with the ACSeg obtained cell masks, i.e. single cell per tomogram. Fig. 1 illustrates the complete workflow of our proposed method. Once the tomograms were masked to get single cells, we performed Sobel-based high pass filter preprocessing and resizing (Materials and Methods 2.4 and 2.5). Next, the tomograms along with ground truth masks for the nucleus and mitochondria were sliced into 3D 64-voxel thick slices along all three axes. The 3D slices had an overlap of 32 voxels along the thickness. This generated a total of 63 3D slices from each tomogram (Fig. 1A). The 3D U-Net model was trained (Materials and Methods 2.6) using either BCE loss or a combination of BCE and Robust Dice loss (Fig. 1B). The details of the loss function and other architectural parameters for the model are mentioned in Materials and Methods 2.7. For prediction on a new cell, the cell was similarly divided into 3D slices, and the probabilistic predictions were merged into a single complete mask by averaging the probability values for each voxel independently (Fig. 1C, Materials and Methods 2.8).

**Fig. 1:**
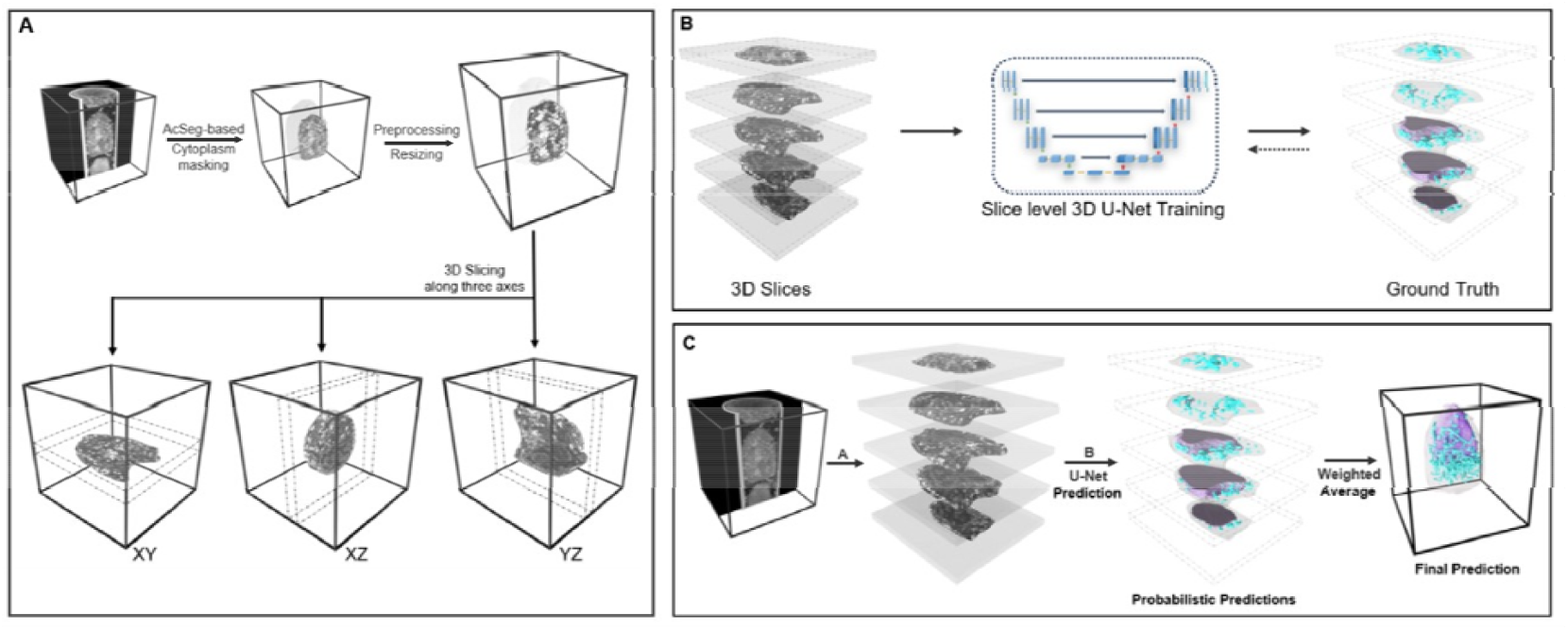
Auto-segmentation workflow for mitochondria and nucleus. (A) 3D slicing: Workflow showing the cell masking using ACSeg, Preprocessing and Resizing, and 3D slicing along the three axes. (B) Training: 3D slices of tomogram and ground truth are used for training the 3D U-Net model. (C) Testing: Test tomogram is sliced into 3D slices as shown in (A). Mitochondria and nucleus masks are predicted using the model trained in (B), and finally, all the predicted masks are merged to generate final whole-cell mitochondria and nucleus masks.

### 3.2 Preprocessing of Raw tomograms

The detection of mitochondria in raw SXT tomograms can be challenging, firstly, due to overlaps in intensity values of different organelle regions (Fig. 2A) and secondly, mitochondria exhibit very complex, tubular, dynamic architecture with varying widths, lengths and shapes (Fig. 2B). Therefore, preprocessing methods that can highlight mitochondria and enhance their boundary are essential for auto-detection and segmentation. The nucleus on the other hand, can easily be detected due to much simpler surface morphology.

**Fig. 2:**
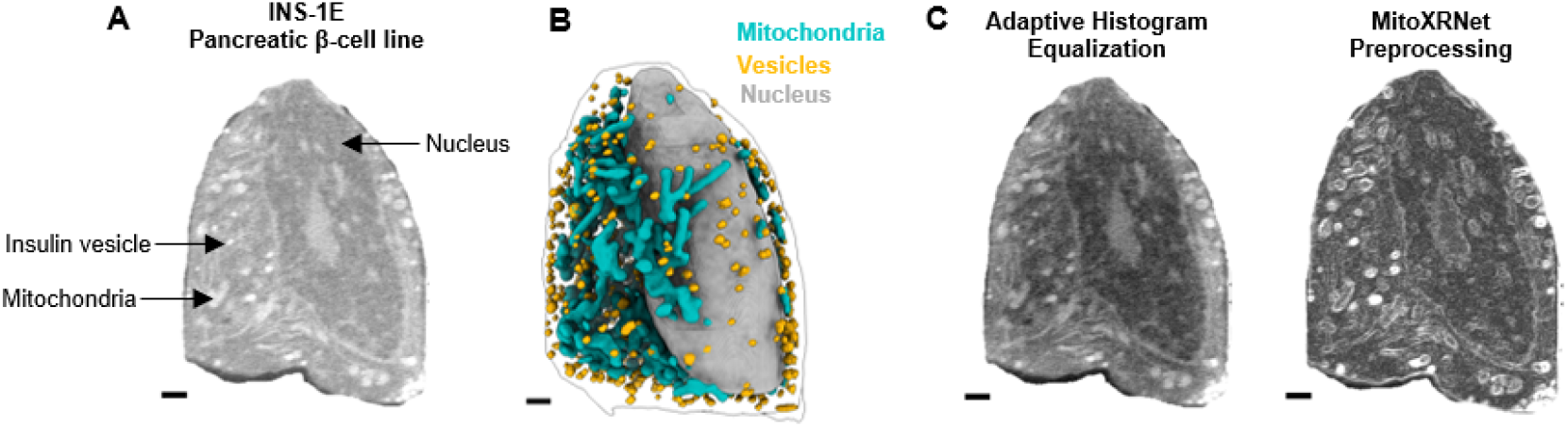
Tomogram preprocessing. (A) 2D slice of a raw tomogram (Image slightly enhanced for visualization). (B) Manual segmentation of the tomogram in (A). (C) Adaptive Histogram Equalization (AHE) and Sobel-based high pass filtering with gaussian smoothing and normalization of the slice shown in (A). Scale bars = 1µm

There are numerous techniques to enhance the contrast and edges of objects in images (Chethan et al., 2019; Ganesan & Sajiv, 2017; Jia et al., 2015; Zhang et al., 2009). MitoXRNet utilizes nonlinear Sobel-based high-pass enhancement with Gaussian smoothing and normalization enhancing the boundary of the mitochondria in the tomogram. Detailed methodology is provided in Materials and Methods 2.4. Fig. 2C shows ortho slices of the tomogram processed with Adaptive Histogram Equalization (AHE) as compared to our proposed Sobel-based high-pass filter. The mitochondrial boundaries are significantly enhanced and much more easily detectable with the proposed filter. We thus used this preprocessing technique for further downstream training and testing.

### 3.3 Comparing MitoXRNet with existing methods

We had a total of 55 tomograms from the first dataset, out of which same three tomograms as done by (Li et al., 2022) were kept in the test set. The rest of the tomograms were divided into training (49) and validation (3) sets. Previous work done on segmenting mitochondria and nuclei from SXT tomograms used 2D slices for training the model (Li et al., 2022). Although the 2D slicing approach gave decent quantitative results (Table 1), the qualitative results were very poor as shown in Fig. 3. The reason being that 2D slices do not preserve the full volumetric context of organelles and subcellar structures. To provide volumetric context and preserve the spatial continuity of mitochondria morphology, we used 3D slices, as described in the previous workflow section.

**Table 1.** Dice score (in %) on in-domain test set. The table compares the dice scores from (Li et al., 2022) (recomputed), nnU-Net (Isensee et al., 2021) and MitoXRNet. MitoXRNet was trained with two separate loss functions. The threshold value represents the cut-off value for converting probabilistic masks to binary masks.

| Cell ID | Li et al. |  | nnU-Net |  | MitoXRNet<br>(1.4 M parameters) |  |  |  |
| --- | --- | --- | --- | --- | --- | --- | --- | --- |
|  |  |  |  |  | BCE Loss<br>(threshold = 0.5) |  | Combined Loss<br>(threshold = 0.6) |  |
|  | Mito | Nucleus | Mito | Nucleus | Mito | Nucleus | Mito | Nucleus |
| 766_8 | 70.4 | 93.92 | 77.65 | 96.14 | 77.72 | 97.20 | 73.70 | 94.85 |
| 784_5 | 58.05 | 92.97 | 67.16 | 94.72 | 77.70 | 96.44 | 75.26 | 95.03 |
| 842_17 | 64.36 | 89.48 | 75.41 | 90.25 | 66.06 | 89.99 | 71.82 | 89.64 |
| Average | 64.27 | 92.12 | 73.41 | 93.70 | **73.83** | **94.54** | 73.59 | 93.17 |
| Std | 5.04 | 1.91 | 4.51 | 2.51 | 5.49 | 3.23 | **1.41** | 2.50 |

Footnote: BCE = Binary Cross Entropy

**Fig. 3:**
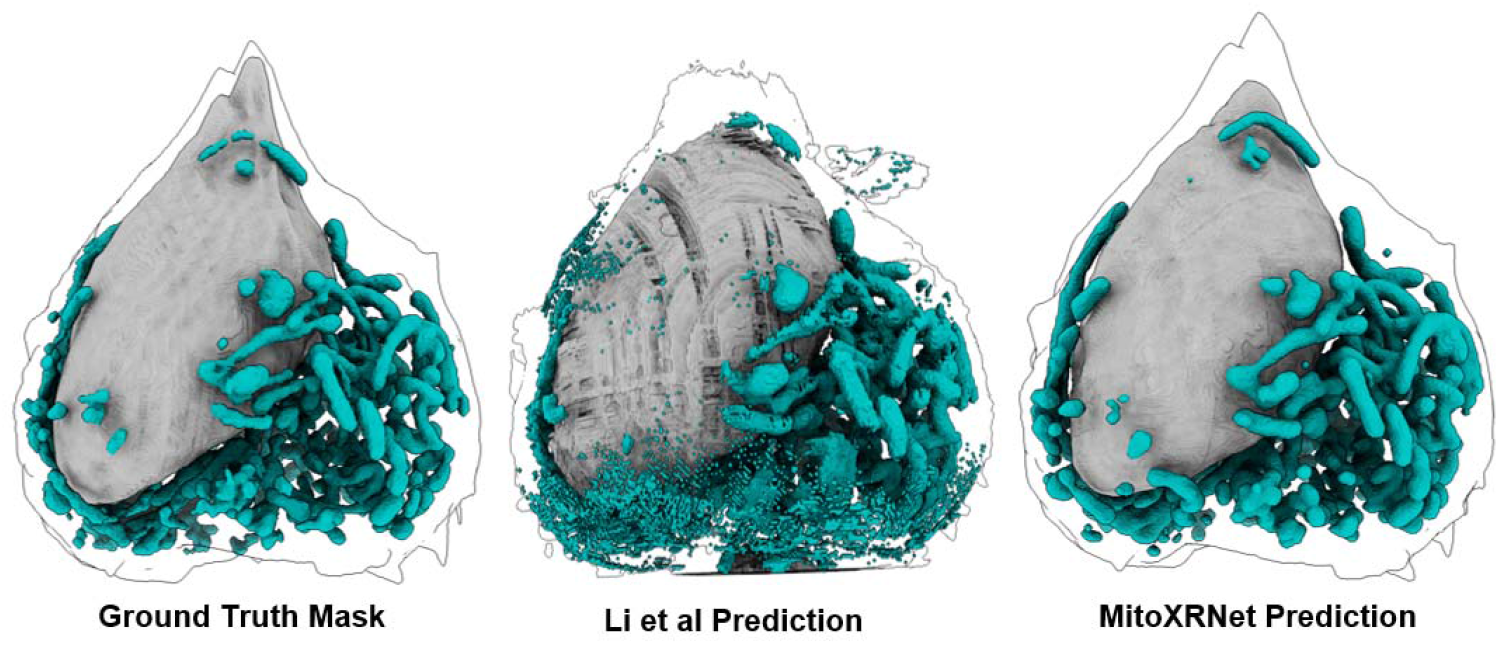
Auto-segmentation comparison: Manual segmentation (Left), Predicted masks of mitochondria and nucleus from (Li et al., 2022) (Middle) and MitoXRNet (Right). Illustrations generate using ChimeraX v1.9 (Goddard et al., 2017)

Initially, we trained a significantly small network with only 1.4 million parameters. The model was trained on two different losses separately: one with BCE Loss and another with a combination of robust Dice Loss and BCE Loss. We saw that the addition of robust dice loss provided stability to overall loss by reducing the deviation in DICE score values (Table 1) and is better where class imbalance is extreme. The proposed approach of using 3D slicing along with preprocessing gives better quantitative (Table 1) and qualitative (Fig. 3) results.

The nnU-Net model auto-configured to a 31.2M parameter full-resolution 3D U-Net model that uses 3D blocks, rather than 3D slices. From (Table 1), we can see that the average Dice score on in-domain test cells is highest for MitoXRNet, outperforming (Li et al., 2022) and nnU-Net (Isensee et al., 2021). Although MitoXRNet achieved the highest dice score for mitochondria with BCE loss (73.83%), the standard deviation is also high (±5.49%), whereas, with combined loss, even though the average value is slightly reduced (73.59%), the deviation decreases significantly (±1.41%).

### 3.4 Comparing manual and auto-segmented masks

To analyze the quality of our predictions and further investigate the reason for still missing the significant amount of Dice score, we compared MitoXRNet segmentations with the ground truth. Fig. 4 shows three example regions from in-domain test set cell (cell ID: 842_17) where the prediction and ground truth did not match. In Fig. 4A, we can see that the ground truth is overly segmented to the extent that it includes surrounding vesicles as part of the mitochondrial network whereas MitoXRNet successfully segments only the actual mitochondrial volume. In Fig. 4B, C we can see the same trend of wrongly segmented vesicles as mitochondria in the ground truth which our model correctly distinguishes and predicts only the true mitochondria. Our analysis shows that the errors on ground truth masks increase false negatives fraction, reducing the overall Dice score.

**Fig. 4:**
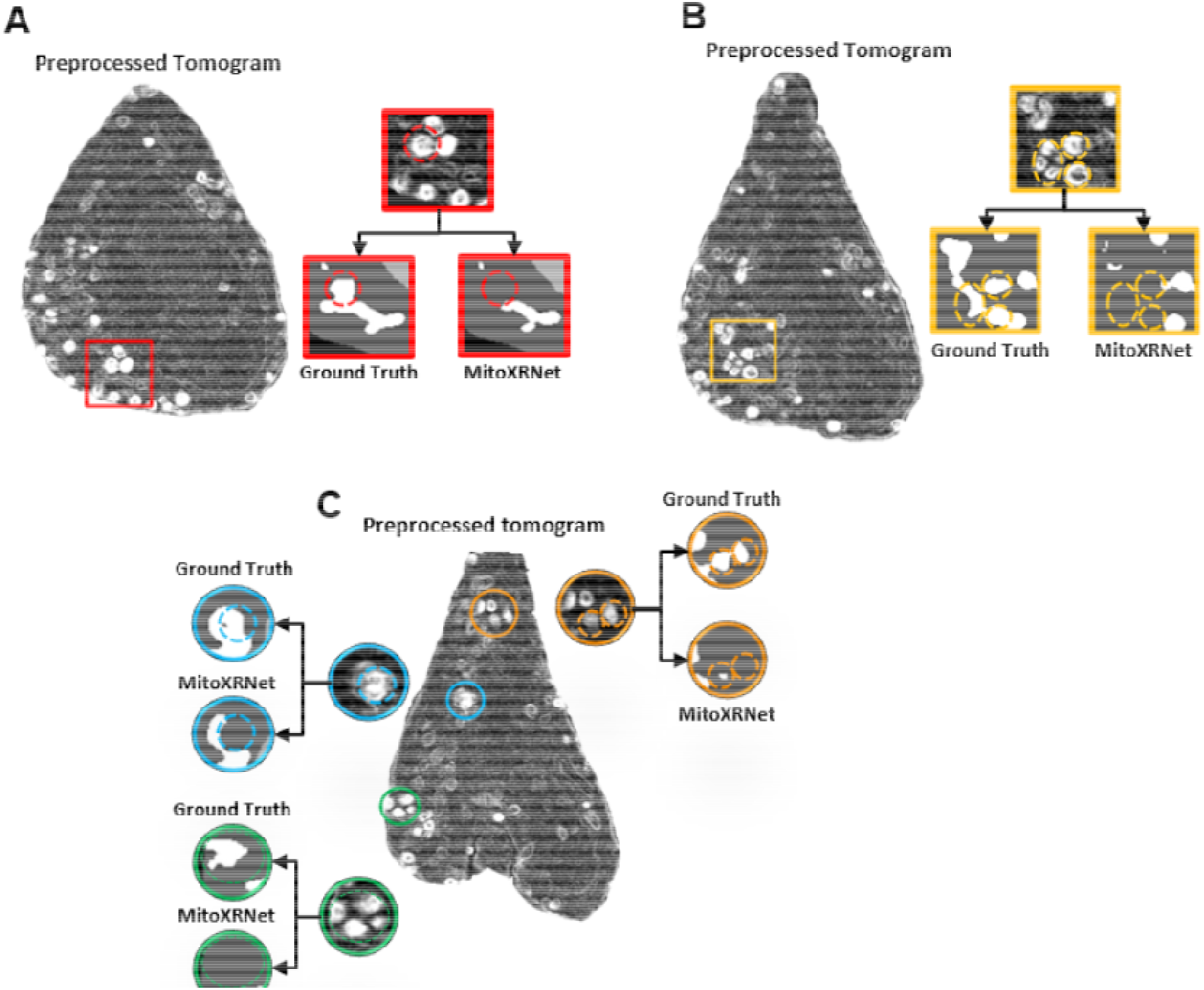
Comparison of predicted and ground-truth masks. (A), (B), (C) Showing three example regions in cell contrast enhanced 842_17 cell, where the ground truth mask of mitochondria includes vesicles, but the predicted mask avoids vesicle region.

### 3.5 Evaluation of unseen out-of-domain test set

We also evaluated the performance of MitoXRNet on an independent unseen out-of-domain test dataset with 44 tomograms. These tomograms were collected to study the effect of various drugs on insulin vesicle trafficking and maturity in pancreatic β-cells (Deshmukh et al., 2025). The details of this dataset are mentioned in Materials and Methods 2.2. Although the 3D U-Net model with 1.4 million parameters performed well on the in-domain dataset, its generalizability appeared limited when evaluated on out-of-domain data. The initial model with 1.4M parameters only achieved 64.63±10.2% with BCE loss, 60.95±15.33% with combined loss, whereas nnU-Net scored 68.96±16.15% average dice score.

So, we trained a larger variant of MitoXRNet with 22.6 million parameters, but still smaller than 31.2 million parameters nnU-Net. When increasing the depth of the model, we saw the same level of performance on the in-domain test set (73.16±3.74%) but significantly improved generalization on out-of-domain dataset (69.28±10.74%.). It did not only perform better than nnU-Net on average dice but also improved on the variation (from ±16.15% to ±10.74%).

### 3.6 MitoXRNet-based automated segmentation reveals mitochondrial morphology changes in pancreatic β-cells

The subset of tomograms of pancreatic β-cells (INS-1E) treated with various secretory stimuli (Deshmukh et al., 2025) were selected for mitochondrial morphology analysis post automated segmentation. These include cells with no stimulation (NS), cells treated with 25mM glucose for 30 min (Glu 30min), and cells co-treated for 30 min with 25mM glucose and 100nM of GLKA-50, a glucokinase activator (Glu+GKA 30min) or 10nM gastric inhibitory polypeptide, an GIPR agonist (Glu+GIP 30 min). After segmentation, whole mitochondrial networks were divided into individual mitochondrial masks, allowing precise quantification of parameters such as mitochondrial number, individual mitochondrial volume, and voxel intensity.

#### 3.6.1 Individual Mitochondrial Volume and Count

Under basal (no stimulation) conditions, mitochondria displayed a wide volume distribution, whereas stimulation with 25 mM glucose for 30 min caused a significant increase in individual mitochondrial volumes (Fig. 5A), indicating structural remodeling under elevated metabolic demand. We observed that very large mitochondria underwent fragmentation, leading to an increased volume and a higher fraction of medium-sized mitochondria (0.2 – 0.6 µm^3^) (Fig. 5D-E). This observation is consistent with previous reports showing that high glucose induces mitochondrial fragmentation or mitophagy in β-cells (Jhun et al., 2013; Serikbaeva et al., 2022; Tseng et al., 2024). In our dataset, the early response appears to be characterized by fragmentation of enlarged mitochondria accompanied by a transient increase in mitochondrial volume.

**Fig 5:**
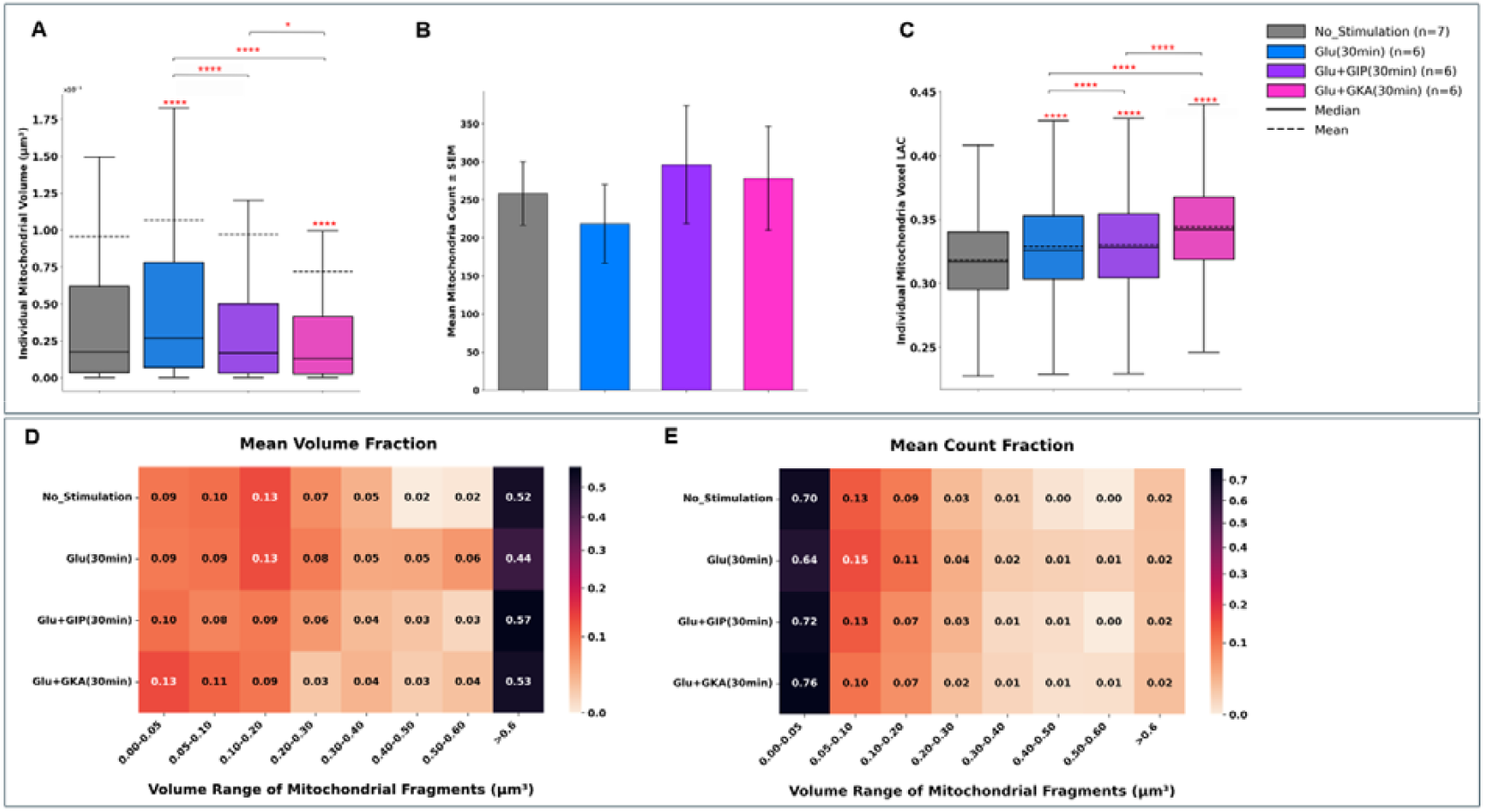
Effects of secretory stimuli on mitochondrial morphology and molecular density in INS1e β-cells. (A) Boxplot of individual mitochondrial volumes (µm^3^) across different secretory stimuli. Solid lines represent median values, and dashed lines denote mean values for each condition. (B) Bar graph of mean mitochondrial count ± SEM reveals distinct effects of each stimulus on mitochondrial abundance, reflecting condition-dependent differences in mitochondrial fragmentation and biogenesis. (C) Boxplot of voxel-level linear absorption coefficient (LAC) values for mitochondrial masks. (D-E) Heatmap of mean volume fraction and mean count fraction at different mitochondrial volume ranges (µm^3^) for each condition, illustrating redistribution of mitochondrial fragments across size classes. Total of 6563 mitochondria across 25 cells were analyzed (No Stimulation: 1,807, Glu 30 min: 1,310, Glu + GIP 30 min: 1,776 and Glu + GKA 30 min: 1,670). *p<0.05; **p<0.01; ***p<0.001, ****p<0.0001 as calculated by Kruskal–Wallis test with post hoc Dunn’s correction in panels (A–C).

Co-treatment with GKA or GIP for 30 min significantly reduced the glucose-induced changes in individual mitochondria volumes (Fig. 5A), but increased the mitochondria count per cell (Fig. 5B), indicated by a shift towards smaller, more numerous mitochondrial units (Fig. 5E). This pattern suggests enhanced mitochondrial turnover or adaptive remodeling rather than maladaptive fission. Such remodeling may contribute to β-cell survival, consistent with the pro-survival actions of GIP reported previously (Kim et al., 2011). The glucose□+□GKA treatment showed the high mitochondrial count compared to glucose (30 min), suggesting that glucokinase activation further drives mitochondrial fission. Heatmaps of mitochondrial volume and count fractions (Fig. 5D–E) confirmed a pronounced redistribution toward the smallest volume range (0.00–0.05 µm^3^), particularly in the glucose + GKA condition, indicating enhanced mitochondrial fragmentation or renewal compared to glucose alone. To our knowledge, this is the first report demonstrating such coordinated mitochondrial count and volume shifts in β-cells following GIP or GKA stimulation.

Overall, our data suggests that GIP and GKA bias mitochondrial remodeling away from large, swollen fragments toward smaller, high-count and possibly more dynamic fragments.

#### 3.6.2 Mitochondrial Voxel intensity

The LAC, or voxel intensity, in SXT tomograms serves as a proxy for biomolecular density and has been thought to reflect the level of metabolic activity (Deshmukh et al., 2025; Loconte et al., 2022; Parkinson et al., 2008; White et al., 2020). Our analysis shows that glucose stimulation alone elevates mitochondrial voxel LAC relative to basal conditions, suggesting increased respiratory activity and ATP synthesis (Maechler et al., 2006). Treatments combining GIP or GKA with glucose further increased the LAC. The increased LAC is consistent with higher oxidative phosphorylation flux, but whether this is adaptive or precedes stress-induced fragmentation requires further investigation.

## 4 Discussion

This study presents a robust and efficient methodology for the auto segmentation of mitochondria and nucleus from Soft X-ray tomograms. We demonstrated that appropriate preprocessing can significantly enhance mitochondrial representation in tomograms thus improving the model’s ability to recognize structural variations. Furthermore, incorporating volumetric context by training on overlapping 3D slices along all three axes rather than 2D slices can substantially improve segmentation quality and accuracy. Using Sobel-based preprocessing, MitoXRNet achieved better segmentation compared to both the 2D segmentation approach by (Li et al., 2022) and the self-configuring nnU-Net (Isensee et al., 2021), despite having only 1.4M parameters for in-domain data and 22.6M parameters for out-of-domain datasets.

In one of the in-domain test cells, we identified multiple regions where vesicles were incorrectly labeled as mitochondria in the ground truth. Such mislabeling increases the contribution of false negatives and lowers the Dice score, indicating that the model’s actual performance is likely to be better than what is indicated by current dice score value. Additionally, we observed that a few insulin vesicles were occasionally predicted as mitochondria due to their morphological similarity with small mitochondria, leading to increase in false positives. To address these errors and further improve the segmentation accuracy, future work could explore boundary-aware loss functions and advanced architectures. Moreover, co-predicting multiple organelle masks could improve the segmentation accuracy for each organelle. However, vesicle segmentation remains challenging due to extreme class imbalance relative to mitochondria and nucleus. Addressing this will require adaptation of adequate preprocessing techniques and loss functions.

We also investigate the morphological changes in mitochondrial volume, count and metabolic activity on treatment of pancreatic beta-cells with glucose with and without secretory stimuli. These stimuli have been studied for their effect on insulin vesicle maturation in the recent study (Deshmukh et al., 2025), but their effect on mitochondria remodeling remains an open question, due to lack of mitochondrial masks. Although the literature contains comprehensive reviews on β-cell mitochondrial bioenergetics, dynamics, and morphology (Supale et al. 2012; Stiles and Shirihai 2012; Diane et al. 2022; Rivera Nieves et al. 2024) 2024) there remains a paucity of data reporting the effect of incretin stimulation (GIP) or glucokinase activation (GKA) on mitochondrial morphology in β-cells. Thus, our findings fill an important gap, implying that these pharmacological modulations not only alter mitochondrial number and size but also internal density properties.

Overall, MitoXRNet establishes a scalable and accurate framework for 3D segmentation of organelles in Soft X-ray tomograms. This work lays the foundation for systematic studies of cellular architecture and its alterations under various physiological and pathological conditions.

## Acknowledgments

The work was funded by Chellaram Diabetes Research Centre (CDRC). We also thank IIT Roorkee for providing additional financial support for computing resources. We thank Valentina Loconte for her contribution to the data collection efforts. We thank Riva Verma for help in designing figures.

## Author contributions

Conceptualization, J.S.; Data curation, A.D. and K.L.W.; Formal analysis, A.Y. and A.S.; Funding acquisition, J.S.; Methodology, A.Y., P.B., A.B. and A.S.; Writing-original draft, A.Y.; Writing-review & editing, A.D, J.S., K.L.W.

